# REDCRAFT: A Computational Platform Using Residual Dipolar Coupling NMR Data for Determining Structures of Perdeuterated Proteins Without NOEs

**DOI:** 10.1101/2020.06.17.156638

**Authors:** Casey A. Cole, Nourhan S. Daigham, Gaohua Liu, Gaetano T. Montelione, Homayoun Valafar

## Abstract

Nuclear Magnetic Resonance (NMR) spectroscopy is one of the two primary experimental means of characterizing macromolecular structures, including protein structures. Structure determination by NMR spectroscopy has traditionally relied heavily on distance restraints derived from nuclear Overhauser effect (NOE) measurements. While structure determination of proteins from NOE-based restraints is well understood and broadly used, structure determination by NOEs imposes increasing quantity of data for analysis, increased cost of structure determination and is less available in the study of perdeuterated proteins. In the recent decade, Residual Dipolar Couplings (RDCs) have been investigated as an alternative source of data for structural elucidation of proteins by NMR. Several methods have been reported that utilize RDCs in addition to NOEs, and a few utilize RDC data alone. While these methods have individually demonstrated some successes, none of these methods have exposed the full potential of protein structure determination from RDCs. To date, structure determination of proteins from RDCs is limited to small proteins (less than 8.5 kDa) using RDC data from many alignment media (>3) that cannot be collected from larger proteins. Here we present the latest version of the REDCRAFT software package designed for structure determination of proteins from RDC data alone. We have demonstrated the success of REDCRAFT in structure determination of proteins ranging in size from 50 to 145 residues using experimentally collected data and large proteins (145 to 573 residues) using simulated RDC data that can be collected from perdeuterated proteins. Finally, we demonstrate the accuracy of structure determination of REDCRAFT from RDCs alone in application to the structurally novel PF.2048 protein. The RDC-based structure of PF.2048 exhibited 1.0 Å of BB-RMSD with respect to the NOE-based structure by only using a small amount of backbone RDCs (∼3 restraints per residue) compared to what is required by other approaches.

**Author Summary:** Residual Dipolar Couplings have the potential to reduce the cost and the time needed to characterize protein structures. In addition, RDC data have been demonstrated to concurrently elucidate structure of proteins, perform assignment of resonances, and be used in characterization of the internal dynamics of proteins. Given all the advantages associated with the study of proteins from RDC data, based on the statistics provided by the Protein Databank (PDB), surprisingly the only 124 proteins (out of nearly 150,000 proteins) have utilized RDCs as part of their structure determination. Even a smaller subset of these proteins (approximately 7) have utilized RDCs as the primary source of data for structure determination. The impeding factor in the use of RDCs is the challenging computational and analytical aspects of this source of data. In this report, we demonstrate the success of the REDCRAFT software package in structure determination of proteins using RDC data that can be collected from small and large proteins in a routine fashion. REDCRAFT accomplishes the challenging task of structure determination from RDCs by introducing a unique search and optimization technique that is both robust and computationally tractable. Structure determination from routinely collectable RDC data using REDCRAFT can lead to faster and cheaper study of larger and more complex proteins by NMR spectroscopy in solution state.

## Introduction

Nuclear Magnetic Resonance Spectroscopy is a well-recognized and utilized approach to structure determination of macromolecules including proteins. NMR spectroscopy has contributed to structural characterization of nearly 11,649 proteins based on statistics reported by the Protein DataBank(1-3) (PDB). Although NMR studies may in general be more time consuming and costly than X-ray crystallography, they provide the unique benefit of observing macromolecules in their native aqueous state, which provide a better understanding of molecular interactions and internal dynamics at various timescales and resolutions.

Despite the changes that NMR spectroscopy has observed over the years, the methodology in analysis of NMR data has made relatively little progress. Nearly all methods of NMR data analysis rely on a combination of Simulated Annealing(4, 5), Gradient Descent(4, 5), and Monte Carlo sampling(4, 5) to guide their protein structure calculations in satisfying the experimental constraints. The traditional approaches to characterization of protein structures by NMR spectroscopy have relied heavily on sidechain-sidechain based distance constraints(6), which are limited to a range of 2.5-5Å. The distance constraints obtained by NMR spectroscopy are often augmented with other heterogenous data such as backbone-backbone contacts, scalar couplings, and dihedral restraints. The structure of the target protein is then computed by deploying a combination of constrained Monte Carlo and Gradient Descent optimization routines. This combination of heterogeneous data and optimization techniques with well documented limitations(4, 7) has resulted in an inflated requirement for experimental data. The functional consequence of this mechanism of protein structure determination has manifested itself as inflated data acquisition time and the cost of structure determination, while also functionally limiting the upper boundary in the size of the proteins that can be studied by NMR spectroscopy.

In the recent years, Redisual Dipolar Couplings (RDC) has been recognized as a promising alternative to the traditional Nuclear Overhauser Effect (NOE) constraints. RDCs have distinct advantages over the traditional distance constraints(8-14). Generally, RDC data are more precise, easier to measure, and are capable of providing informative structural and dynamic information. Structure determination based primarily on RDC data requires new approaches that operate in fundamentally different ways from those that use NOE data. This is the primary reason that the legacy programs such as Xplor-NIH(15), CNS(16) or CYANA(17) are not appropriate for de novo structure determination purely by RDC data. Other contemporary methods have been presented(8, 13, 18-25) with a direct focus on characterization of structure from RDC data. While these programs address some of the shortcomings of the traditional approaches, their continued use of the conventional optimization techniques prevent full utilization of the rich information content of the RDC data. Some of these algorithms exhibit a direct or indirect reliance on completeness of the PDB and a thorough sampling of the protein fold-space(23, 24). Others utilize impractical number of RDCs(25-28) (e.g. 4 RDCs per residue collected in 5 alignment media) that cannot be routinely collected, especially on larger and perdeuterated proteins. Furthermore, all the existing RDC-based structure determination approaches deploy search strategies that continue to rely on the traditional optimization techniques such as Levenberg-Marquardt(29) or gradient descent. These approaches work for meticulously clean and complete datasets and therefore lack the needed practical robustness for the analysis of noisy or missing data. Finally, there is no currently existing software that is capable of concurrent structure determination and identification of internal motion in proteins. REDCRAFT illustrates a number of unique advantages, with its most unique feature consisting of a novel search methodology optimally suited for the analysis of RDC data.

Here we demonstrate the extended abilities of REDCRAFT in structure calculation of proteins from Residual Dipolar Couplings (RDCs) that can be collected routinely from small and large proteins. More specifically, we demonstrate the feasibility of structure determination of proteins using only RDCs that can be obtained from perdeuterated proteins, namely backbone {C’-N, N-H^N^, C’-H} in two alignment media. When available, our investigations are based on previously reported experimental RDC data, and when needed, the experimental data are augmented with synthetic data. We have demonstrated successful structure determination by REDCRAFT on eight proteins with a size range of 50 to 573 amino acids. Finally, REDCRAFT has been tested in structure determination of a novel protein, PF2048.1, and the results were validated in comparison to NOE based structures.

## Results

In the following sections we present three sets of results, all of which demonstrate structure determination of proteins from RDCs alone to reduce the overall cost of structure determination. In the first set, we explored the structure determination of all proteins by REDCRAFT, for which sufficient experimental RDC data were deposited into the BMRB database. In each of these exercises, we used substantially smaller set of RDC data than the previously reported studies. In the second set of results, we have investigated the success of REDCRAFT in structure determination of large proteins using synthetically generated RDCs. The structure of these proteins had been previously characterized by distance constraints while including a very small subset of RDCs, therefore establishing the plausibility of RDC collection for these proteins. In the third exercise, we have characterized the structure of the novel protein PF2048.1 by the traditional NOE data and RDC data separately. Using the two structures, we established the accuracy of the RDC based structure from REDCRAFT to the conventional NOE based structure with using only a fraction of the data.

### Protein Structure Calculation Using Experimental RDCs

Table 1 and Figure 1 summarize the results of REDCRAFT structure calculation of proteins using only experimental RDC data. The listed five proteins in this table have been previously studied by NMR spectroscopy, for which enough experimental RDCs have been deposited to BMRB(30). In some instances, RDC data are missing for a large portion of a protein. In such instance, REDCRAFT accommodates fragmented structure determination and therefore the structural comparison to the target structure is reported as a range of BB-RMSD’s calculated for each fragment. The fourth and fifth columns of Table 1 provide a quality of structural fitness to the RDC data as a Q-factor(31). In summary, a Q-factor over the threshold of 0.3 is considered a poorly fit structure, while values less than 0.2 are considered acceptable, and values less than 0.1 are considered exceptionally well-fit structures. The last column of Table 1 indicates the percentage of the data that was utilized by REDCRAFT compared to the number of constraints used previously. In general, as seen in Table 1, the obtained structures were less than 2Å from the target structures with sufficiently low Q-factors (indicating a reliable structure), while reducing the total data requirement by as much as 90%. In the following paragraphs, additional detailed results for each protein are discussed.

**Table 1.** Results for structure calculation using experimental RDCs is summarized.

| Target Name | # Res. | BB-RMSD to NMR Structure (Å) | BB-RMSD to X-Ray Structure (Å) | Q Factor of Prev. Solved Structure | Q Factor of REDCRAFT Structure | % of Used Data |
| --- | --- | --- | --- | --- | --- | --- |
| GB1 | 54 | 1.189 | 1.481 | 0.15, 0.11 | 0.098, 0.12 | 22% |
| GB3 | 54 | 1.9 - 2.5 | 1.3 - 2.2 | 0.034, 0.045 | 0.01 - 0.02, 0.02 - 0.34 | 23% |
| Rubredoxin | 50 | 1.121 | 1.02 | 0.08, 0.52 | 0.33, 0.19 | 41% |
| ChR145 | 145 | 1.4 - 2.3 | N/A | 0.17, 0.18, 0.24 | 0.01 - 0.046, 0.01 - 0.038, 0.01 - 0.026 | 11% |
| SR10 | 145 | 1.1 - 2.4 | 1.6-2.4 | 0.9, 0.68, 0.89 | 0.028 - 0.05, 0.05 - 0.16, 0.06 - 0.32 | 18% |

**Figure 1.**
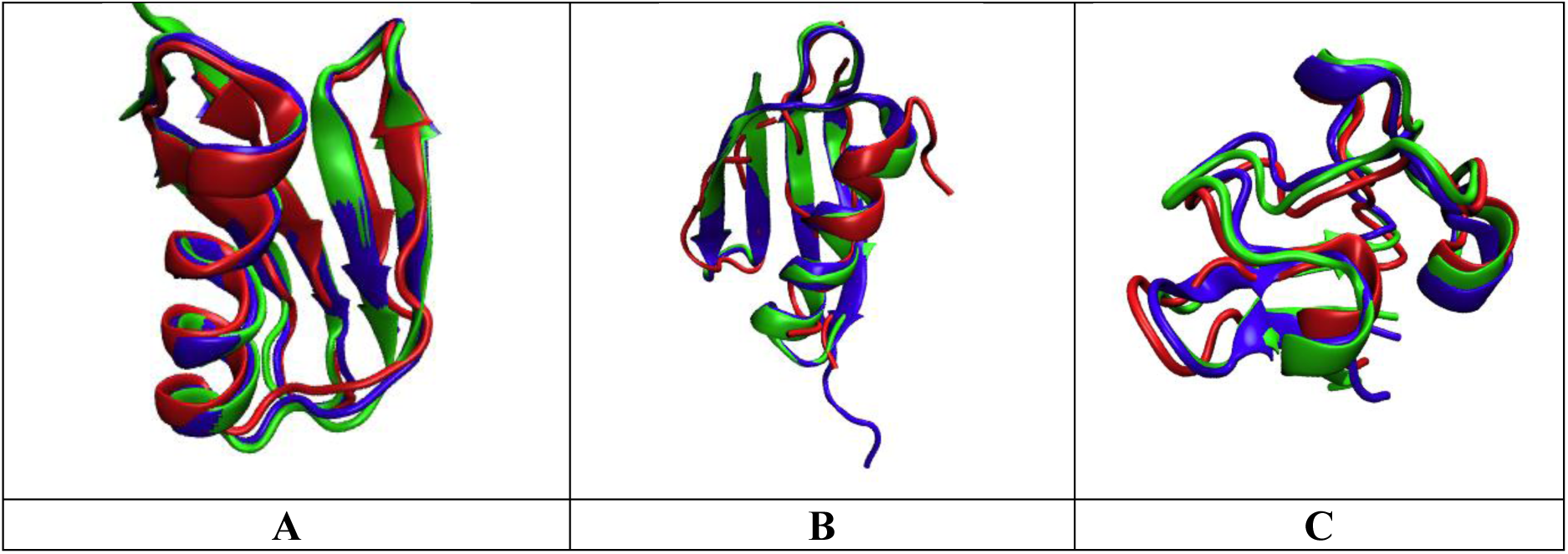
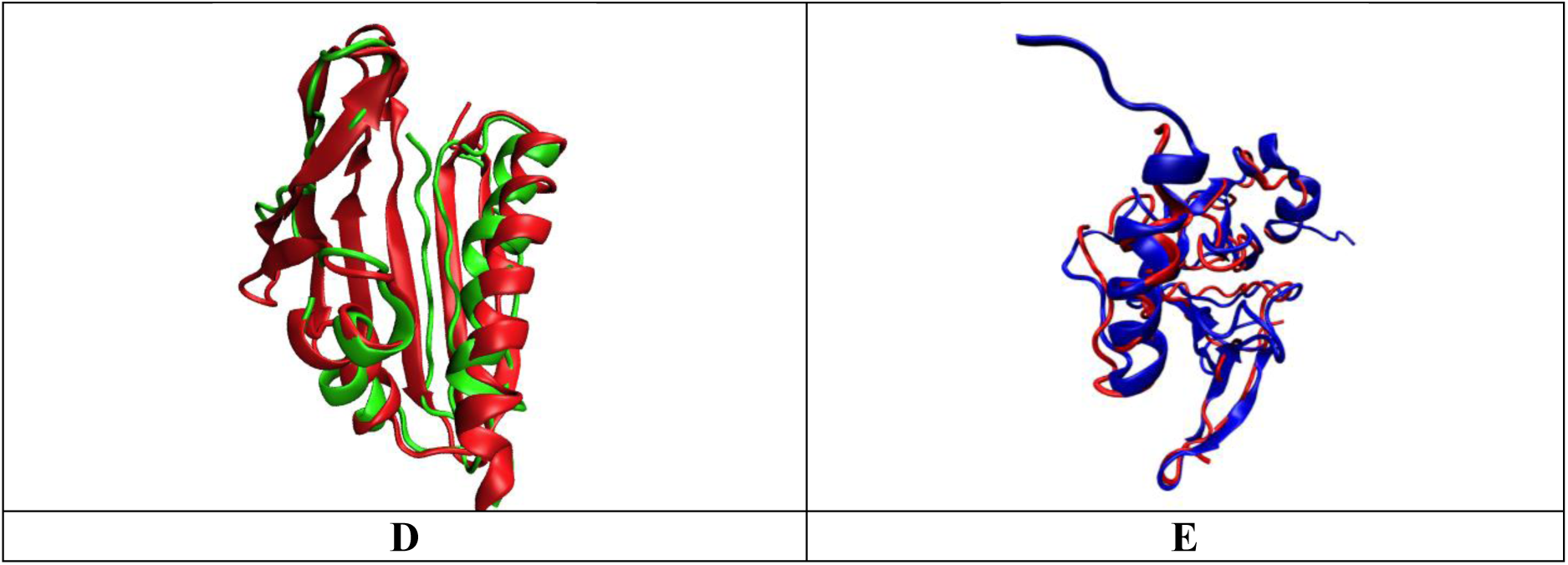
Results of REDCRAFT structure calculation (in red) compared to X-Ray structures (in blue) and, where applicable, traditional NMR Structures (in green) for A) GB1, B) GB3, C) Rubredoxin, D) ChR145, E) SR10.

*GB1 –* The previously calculated NMR structure of GB1 (2PLP) was determined using 769 RDC restraints that included {N-H^N^, N-C’, C_α_-C’, H-C_α_, H-C’, C_α_-C_β_}, 127 long range H^N^-H^N^ RDCs, and 54 Residual Chemical Shift (RCS) restraints from two alignment media(28). In this study, 209 RDC restraints (compared to the total of 950 restraints) were used to obtain a structure less than 1.5Å from both the X-Ray and NMR structures.

*GB3 –* For GB3, the dataset included the following RDCs from: {N-C’, N-H^N^, C_α_-H_α_, C_α_-C’} in five alignment media. Two previous studies used this full set of RDC data to determine a structure of GB3 to within 1Å of the corresponding X-ray structure(32, 33). In a previous REDCRAFT study(34), the structure of GB3 was determined using {N-H^N^, C_α_-H_α_} RDCs in two alignment media. However, the collection of {C_α_-H_α_} RDCs is uncommon due to sample preparation requirement and added complexity in NMR data interpretation. Using these RDCs, REDCRAFT was able to reconstruct the structure to within 0.6-2.4Å of the NMR structure. For the purposes of this study, the RDC data was reduced to the vectors {N-C’, N-H^N^} since they can be collected from perdeuterated protein samples. Utilization of these RDCs are more challenging than the previous set due to their planarity with one another. Using these vectors, we were able to calculate a structure of this protein with BB-RMSD of less than 2.5Å.

*Rubredoxin –* Previously, the structure of Rubredoxin was characterized to within 1.81Å of the X-ray structure using the following RDC vectors: {N-C’, N-H^N^, C’-H, C_α_-H_α_, H^N^-H_α_, H_α_-H^N^} in two alignment media(10). Again, to simulate an RDC set that could be collected from a perdeuterated protein, that experimental RDC set was reduced to {N-H^N^, C’-H} from two alignment media. A BB-RMSD of 1.12Å and 1.02Å were obtained in relation to the NMR and X-ray structures.

*ChR145 –* As part of the original study of ChR145, {N-H^N^, C’-N} RDCs were collected in two alignment media (PEG and Phage) and an additional set of {N-H^N^} RDCs were collected in a third alignment medium (PAG). All RDCs were deposited into the SPINE(35) database. Utilizing only these RDC restraints, REDCRAFT was able to produce structures with BB-RMSDs in the range of 1.4Å −2.3Å. This is then compared to the traditional NMR structure that utilized 2,676 NOEs, 256 dihedral restraints and the 328 RDC restraints.

*SR10 –* The structure of SR10 was obtained by NMR spectroscopy to within 2.0-2.5Å with respect to an X-ray structure. The RDCs available for this protein were 3 sets of {N-H^N^} RDCs. A fragmented study was utilized in this case due to large gaps in the RDC data. The original NOE-based structure utilized 1765 restraints (a mix of RDCs and NOEs) whereas our method only used 320 RDC data.

### Conventional and REDCRAFT Based Structure Determination of PF2048.1

Following the procedures outlined in the Materials and Methods section, two structures were characterized and deposited in the Protein Data Bank(36): one structure that did not include any RDC data that used a total of 2,574 total restricting restraints (PDB_id 6E4J, BMRB_id 30494), and one structure that included RDC (2,534 restricting restraints) (PDB_id 6NS8 and the same BMRB_id 30494). Comparison of these two structures demonstrates the impact of RDC during the last stages of structural refinement following the conventional methods of structure determiantion. Structure quality assessment metrics for these two NMR structures are presented in Supplementary Table S1. Overall, both structures (with and without RDCs) exhibited high quality structures, with excellent structure quality scores. The RDC Q-factors for the two alignment media M1 and M3 are 0.340 ± 0.020 and 0.320 ± 0.031, respectively for the models generated without RDCs, and 0.275 ± 0.015 and 0.280 ± 0.028, respectively, for models generated using RDC data as restraints. The DP scores, assessing how well the models fit to the unassigned NOESY peak list data, are 0.905 and 0.905 for the structures modeled without and with, respectively, RDC data. Molprobity packing scores(37), Richardson backbone dihedral angle analysis(38), and ProCheck(39) backbone and sidechain dihedral angle quality scores for well-defined regions of these models, are also excellent. The backbone RMSD between the medoid models of the ensembles generated with and without RDC data is 0.745 Å. Taken together, this structure quality analysis demonstrates that the experimental NMR structures determined using the conventional approaches are excellent quality, and good reference states for assessing modeling methods using RDC data alone.

The structure of PF2048.1 was determined with only 228 RDCs that consisted of the backbone {C’-H, N-H^N^, C’-N} vectors from the first alignment medium and backbone {N-H} from the second alignment medium. The final REDCRAFT structure exhibited 2.3Å of structural deviation from the target NOE structure before any structural refinement. This structure was then subjected to 20,000 rounds of Powell minimization in Xplor-NIH software package using the same 228 RDC restraints in order to resolve some minor Van Der Waal collisions. The Q-factors before and after minimization for both alignment media are shown in Table 2. It was shown that the Q-factor for M2 was slightly improved. Note that the Q-factor for M1 incurred a slight increase during minimization due to the correction of a Van der Waal collisions in the computed structure. Figure 2 illustrates superimposition of the REDCRAFT computed structure of PF2048.1 (in red) before and after minimization and the NOE structure without RDCs (in blue) and the NOE structure with RDCs (in yellow). The final structure exhibited Q-factors of 0.09 and 0.13 in the two alignment media respectively and a BB-RMSD of less than 1.0 Å with respect to both NOE structures (structures determined with and without RDCs).

**Table 2.**
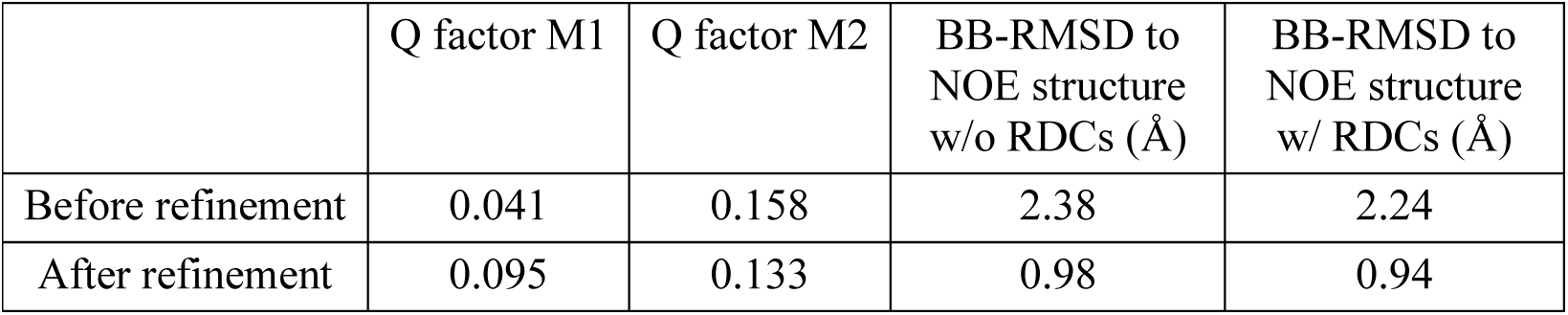
Results from structure calculation of PF2048.1 using 228 RDCs and secondary structure restraints are shown.

|  | Q factor M1 | Q factor M2 | BB-RMSD to<br>NOE structure<br>w/o RDCs (Å) | BB-RMSD to<br>NOE structure<br>w/ RDCs (Å) |
| --- | --- | --- | --- | --- |
| Before refinement | 0.041 | 0.158 | 2.38 | 2.24 |
| After refinement | 0.095 | 0.133 | 0.98 | 0.94 |

**Figure 2.**
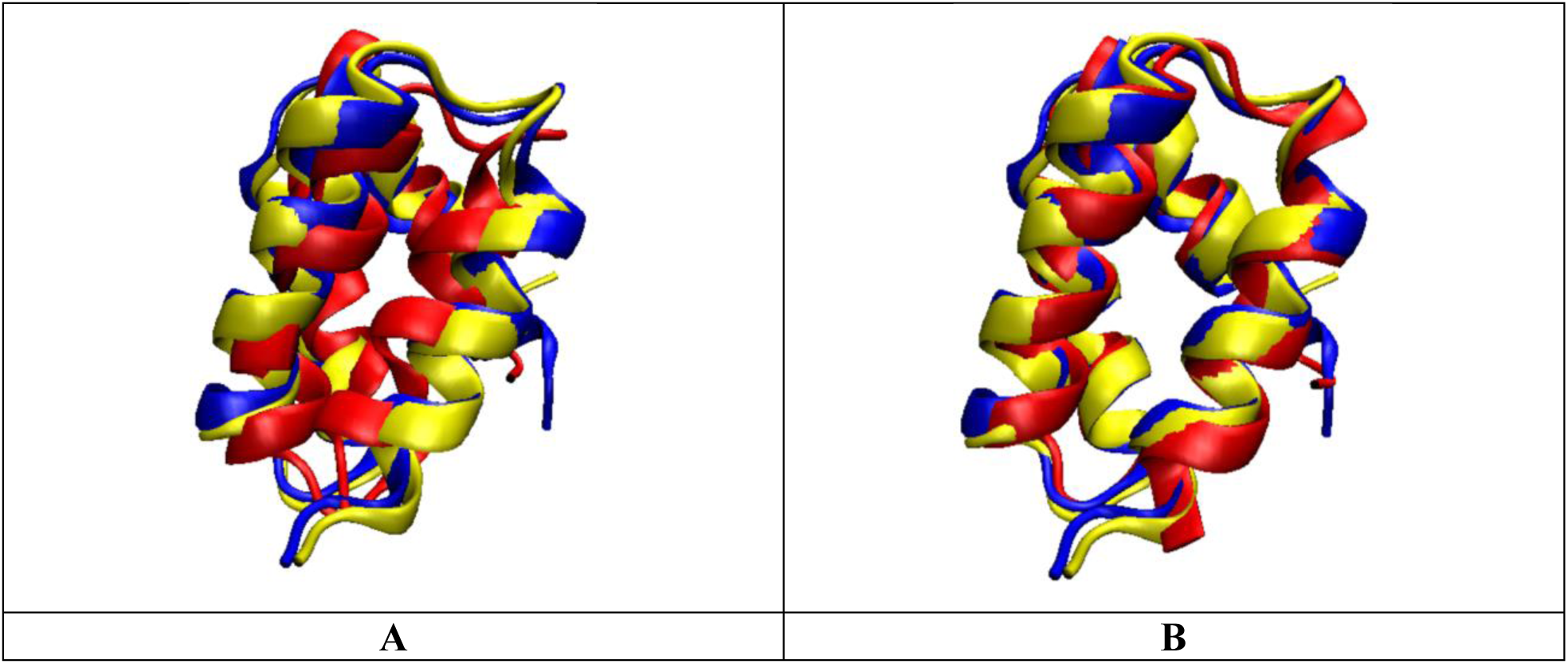
Results for PF2048.1 (in red) A) before minimization and B) after minimization are shown superimposed to the NMR structure without RDCs (in blue) and the NMR structure with RDCs (in yellow).

### Structure Calculation of Large Proteins

The results of structure calculation for large proteins using synthetic RDCs are shown in Table 3 and Figure 3. Although the structure of ChR145 was characterized by REDCRAFT using experimental data (reported in Table 1), here we have repeated the structure determination of this protein with synthetic data to illustrate the possibility of full structure determination (instead of a fragmented study) if adequate RDCs were collected. In this study, ChR145 was characterized in one full continuous segment with a BB-RMSD of 1.45Å with respect to the reference structure. In addition, the resulting structure had excellent Q-factors.

**Table 3.**
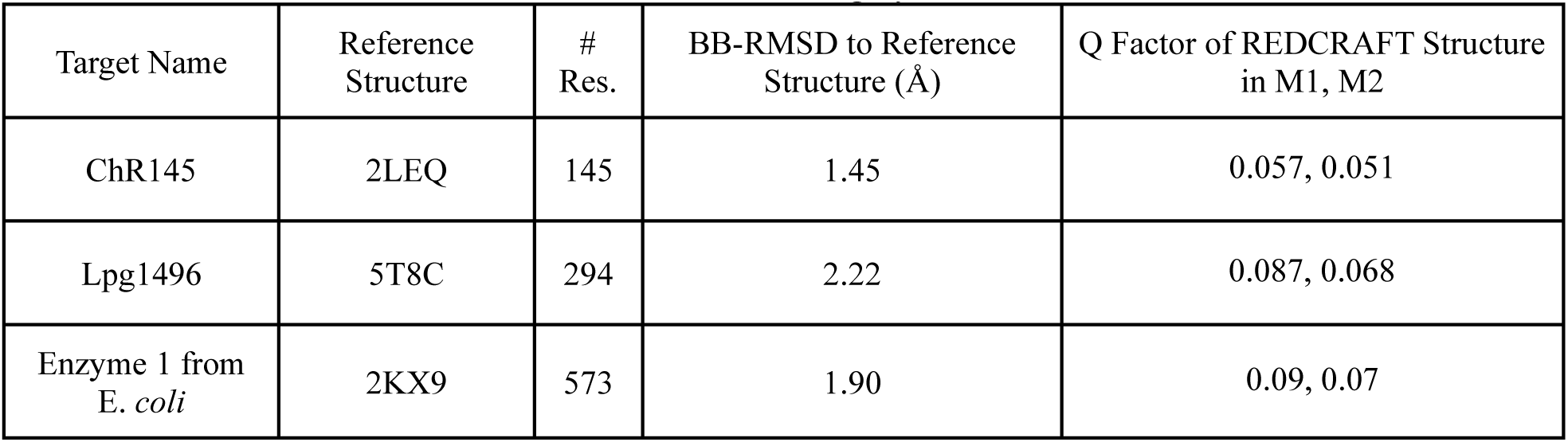
Results for structure calculation using synthetic RDCs is summarized.

| Target Name | Reference Structure | # Res. | BB-RMSD to Reference Structure (Å) | Q Factor of REDCRAFT Structure in M1, M2 |
| --- | --- | --- | --- | --- |
| ChR145 | 2LEQ | 145 | 1.45 | 0.057, 0.051 |
| Lpg1496 | 5T8C | 294 | 2.22 | 0.087, 0.068 |
| Enzyme 1 from *E. coli* | 2KX9 | 573 | 1.90 | 0.09, 0.07 |

**Figure 3.**
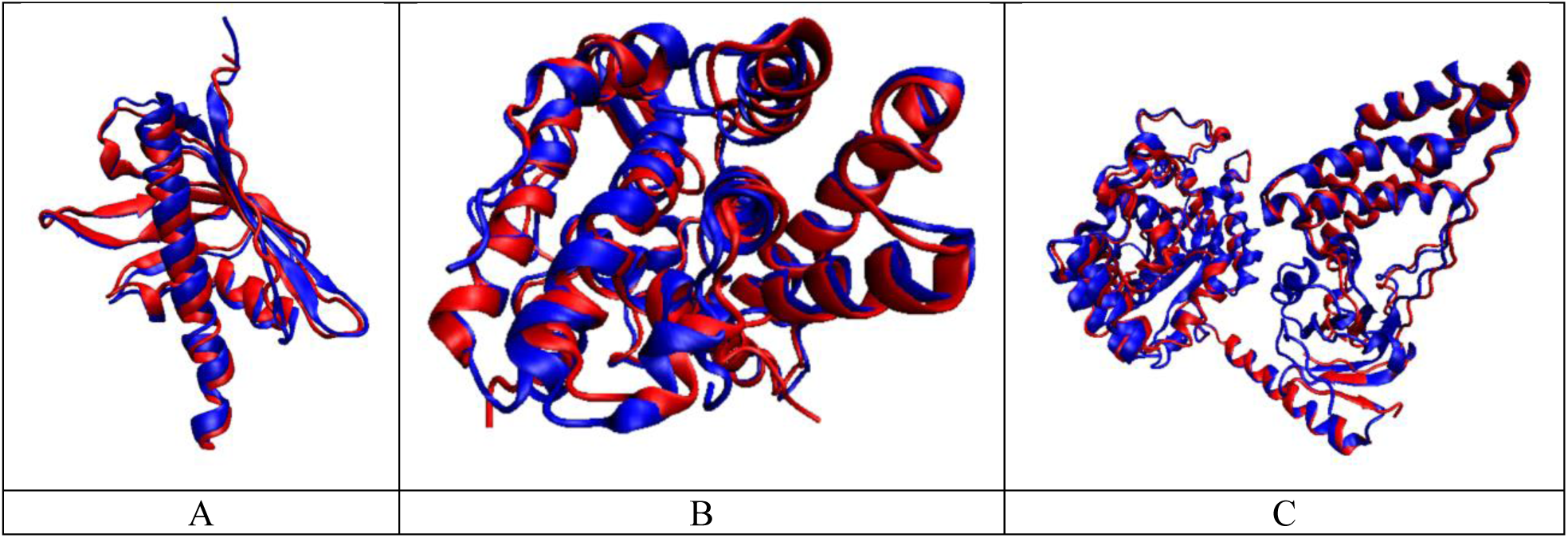
Results of REDCRAFT structure calculation (in red) compared to the reference structure (in blue) A) ChR145, B) Lpg1496 and C) Enzyme 1 from E. *coli*.

In the cases of Lpg1496 and Enzyme 1, fragmented study was performed due to contribution of structural noise discussed in the Materials and Methods section. For instance, in several cases, a single residue’s dihedral angles were in severe violation of the Ramachandran space. In such instances, the structure determination was augmented with short refinement of each fragment followed by their integration using Xplor-NIH. For Lpg1496, the largest contiguous fragment characterized as 138 residues in length displaying a BB-RMSD of 1.73Å. Additional fragments ranged from 50 to 75 residues in length. All fragments reported Q-factors indicative of reliable structure in each alignment medium as well as low BB-RMSDs to the reference structure. The longest fragment for Enzyme 1 was 208 residues, which exhibited a BB-RMSD of 1.78Å. All other fragments ranged from 50 to 100 residues in length. For the fragmented studies, all fragments were aligned to their respective structures and an average BB-RMSD was calculated (shown in the table).

## Discussion

Structure calculation of large proteins from RDCs using REDCRAFT is possible and can have numerous advantages. In this study, we have demonstrated the feasibility of calculating the structure of proteins of varying sizes (50-573 amino acids) from RDCs using REDCRAFT. In addition, due to the novel search mechanism of REDCRAFT, we have demonstrated reliable folding of proteins with as little as 11% of the previously used data. Furthermore, we have shown that RDCs collected on perdeuterated proteins are sufficient for folding large proteins (as large as 573) with high accuracy. This is a significant achievement since in most cases, large proteins must be perdeuterated to be amenable for study by NMR spectroscopy. Using simulated RDCs, we also demonstrated that determination of large proteins is in fact possible using the minimal set of RDCs that can be acquired using perdeuterated samples. Lastly, we show REDCRAFT’s ability to characterize an unknown protein with very little sequence or structural similarity to other proteins.

Structural elucidation of proteins from RDCs using REDCRAFT has other pragmatic advantages. For instance, characterization of protein structure does not have to be restricted to the entire protein. RECRAFT’s approach allows for structural investigation of a fragment of the protein as demonstrated with proteins GB3, ChR145, and SR10. Isolated study of a targeted fragment of a protein will reduce the cost of structure determination and allow for study of larger proteins in a partitioned fashion. Furthermore, the combination of RDCs when analyzed with REDCRAFT, enables concurrent study of structure and dynamics of a protein as presented previously(34, 40, 41), also reducing the cost of such studies.

## Methods

### Residual Dipolar Couplings

Residual Dipolar Couplings (RDCs), an alternate source of data obtainable by NMR spectroscopy, had been observed as early as 1963 in pneumatic solutions(42). The recent reintroduction of RDCs due to the development of alignment media has presented this source of data as a possible substitute to the conventional approach to structure determination by NMR spectroscopy. RDCs have been shown to be valuable for structural characterization of aqueous proteins(10, 12, 43, 44) and challenging proteins(40, 45-49), while enabling simultaneous study of structure and dynamics of proteins(9, 27, 40, 45, 50-53). Because RDCs can be used to characterize the structure of proteins with far less data than the traditional approaches, it presents a viable and cost-effective method of protein structure elucidation.

RDCs arise from the interaction of two magnetically active nuclei in the presence of the external magnetic field of an NMR instrument(54-57). This interaction is normally reduced to zero, due to the isotropic tumbling of molecules in their aqueous environment. The introduction of partial order to the molecular alignment reintroduces dipolar interactions by minutely limiting isotropic tumbling. This partial order can be introduced in numerous ways(58), including inherent magnetic anisotropy susceptibility of molecules(59), incorporation of artificial tags (such as lanthanides) that exhibit magnetic anisotropy(54), or in a liquid crystal aqueous solution(58) as illustrated in Figure 4. The RDC interaction phenomenon has been formulated in different ways(57, 60). In order to harness the computational synergy of RDC data, we utilize the matrix formulation of this interaction as shown in Eq (1). The entity *S* shown in Eq (1) and (2) represents the Saupe order tensor matrix(42, 54, 61) (the ‘order tensor’) that can be described as a 3×3 symmetric and traceless matrix. *D*_*max*_ in Eq (1) is a nucleus-specific collection of constants, *r*_*ij*_ is the separation distance between the two interacting nuclei (in units of Å), and *v*_*ij*_ is the corresponding normalized internuclear vector

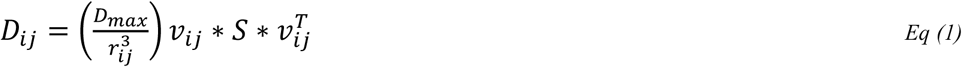

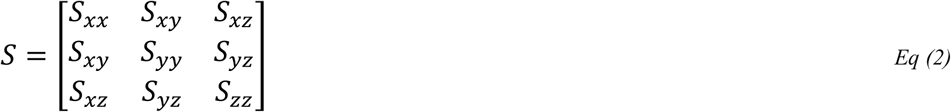

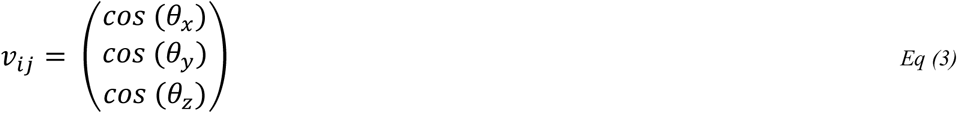

**Figure 4.**
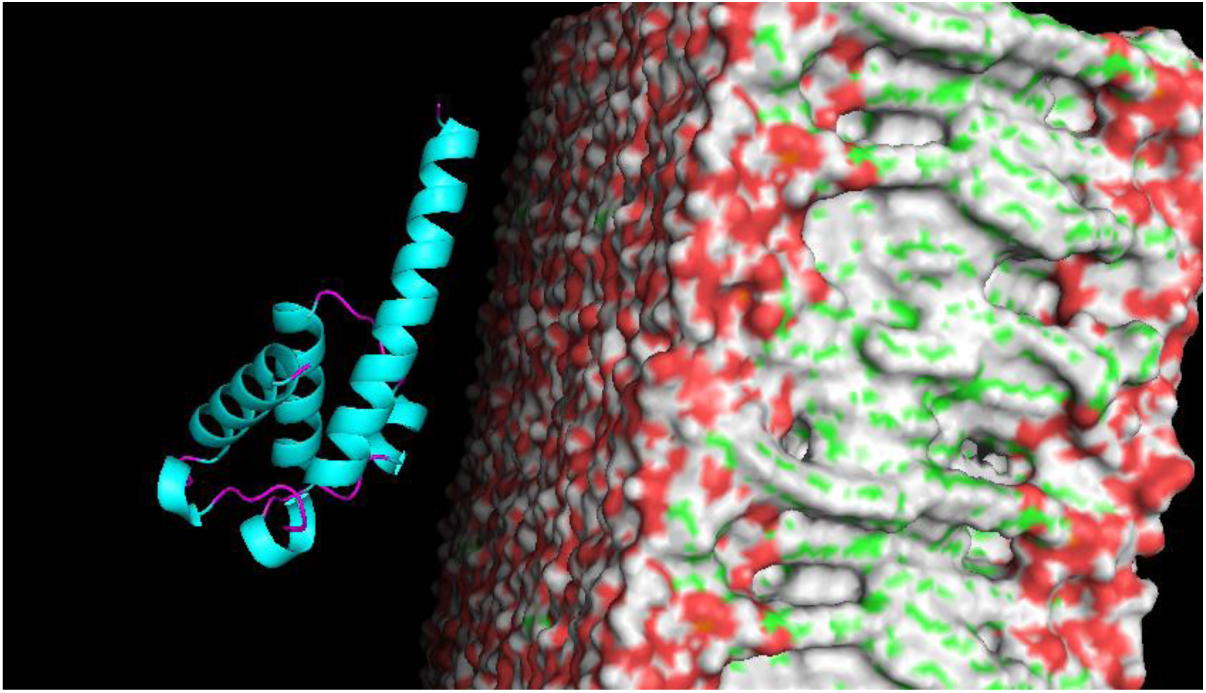
Bicelle crystalline solution is one method of inducing partial alignment.

RDC data have several advantages over the conventional NOE data(8-14). In the interest of brevity, we focus our discussion on the importance of RDC data in high-resolution structure determination. Figure 5 represents the RDC and NOE fitness of 5000 derivative structures as a function of their BB-RMSD to the known structure. These 5000 structures have been derived from a target protein by randomly altering the backbone torsion angles to achieve a continuum of distortions (measured in BB-RMSD’s). Fitness to the experimental data (similar to Q-factor(31)) is calculated and plotted on the vertical axis, while BB-RMSD to the high-resolution structure is plotted on the horizontal axis. This figure illustrates the sensitivity of NOEs and RDCs as reporters of protein structures. Figure 5 suggests that NOEs tend to lose sensitivity as the search approaches the native structure, while RDCs become more sensitive. RDCs can also report molecular motions on timescales ranging from picoseconds to microseconds(62-64), during which many functionally important events occur. Indeed, in the 10 ns – 1 s timescale window, RDCs are the most sensitive of NMR parameters^32^.

**Figure 5.**
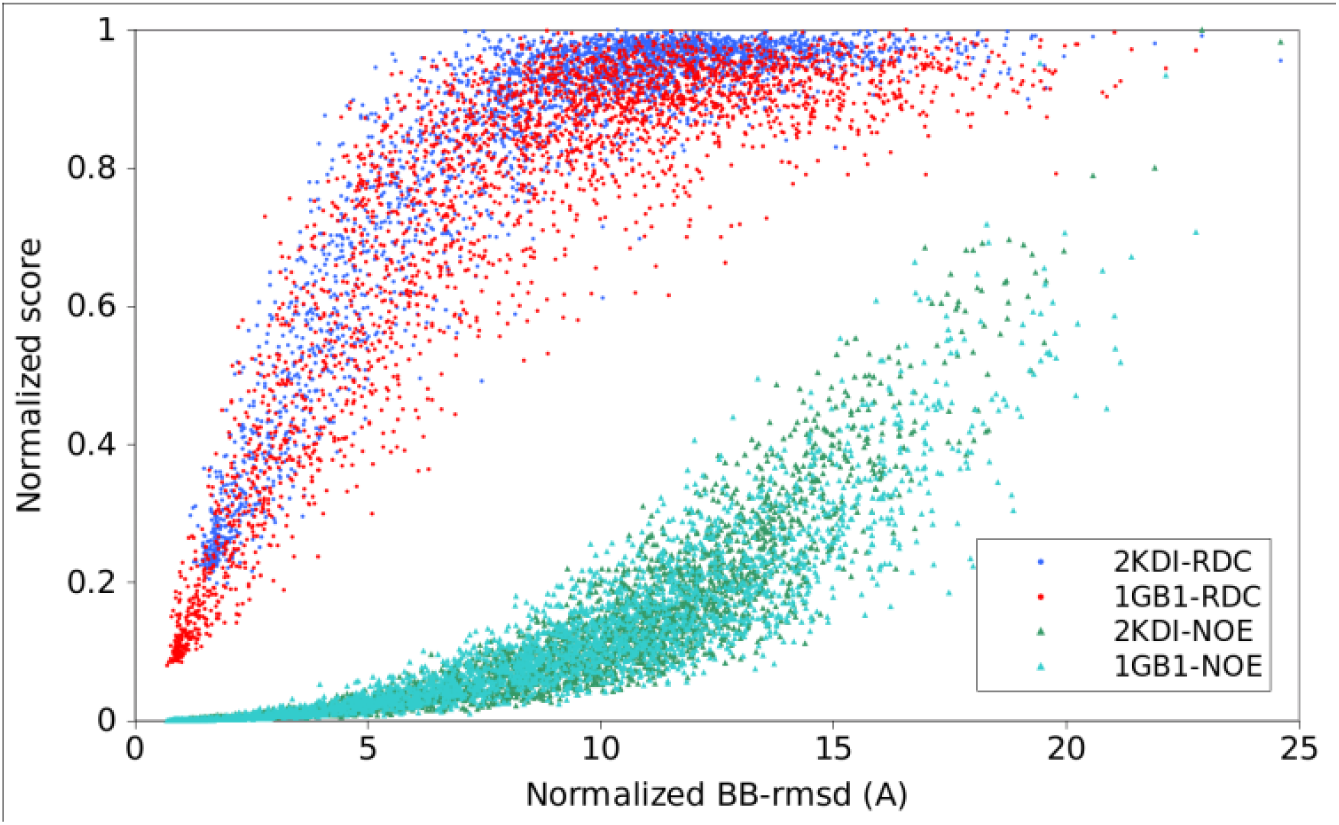
RDC and NOE fitness of 5000 decoy structures generated randomly from a known structure versus their backbone rmsd to the actual structure.

Despite many advantages of RDCs in characterization of protein structures, only a handful of protein structures submitted to the PDB have been determined exclusively by RDC data. Nearly all of the structures determined by RDCs have utilized an excessive number of RDCs than necessary(25-28) or resorted to the use of other experimental data such as NOEs, dihedral restraints and hydrogen bond restraints(21, 65, 66) for successful structure determination. Both of these approaches nullify the advantages of using RDCs (minimal data, reduced cost), and have therefore resulted in the failure to realize the full potential of RDCs.

### REDCRAFT

Realization of the full potential of RDC data has been historically hindered by the lack of appropriate analysis tools. Practically, all of the legacy NMR data analysis software such as Xplor-NIH, CNS, or CYANA have been modified to incorporate RDC data into their analysis. However, the energy landscape that is created by the RDC data is far too complex to be navigated by simplistic optimization routines such as Gradient Descent or Monte Carlo sampling. Therefore, the legacy software can use RDC data only when they are accompanied by a large number of traditional constraints. Structure determination based primarily on RDC data requires new programs that operate in fundamentally different ways than those that use traditional constraints such as NOE data.

In recent years, several computational approaches have been introduced with a focus on structure determination of proteins from RDC data. REDCRAFT is such a program that sets itself apart from other existing software packages in a number of ways. By deploying a novel search mechanism that is significantly different than traditional optimization techniques, REDCRAFT can accomplish the same outcome as other algorithms but with less data. REDCRAFT also presents the ability for simultaneous study of structure and dynamics of proteins, and its predecessor has demonstrated the capability for simultaneous structure elucidation and assignment of data. Using simulated data, the success of REDCRAFT has been demonstrated with as little as two RDCs per residue(34, 67, 68), or one RDC per residue(67, 69) when combined with backbone torsion angle constraints. Meaningful structure determination, based on a subset of RDC data (two or three per residue) obtained from the perdeuterated proteins, is critical in extending the structure determination based on RDCs to large proteins. It is a common practice to deuterate the sample of large proteins, leaving a far smaller selection of RDCs that can be collected. In particular, the set of {C’-H, N-H, C’-N} RDC data can be obtained from deuterated proteins with relative ease. It is therefore of great importance for any RDC-based structure determination technique to be able to characterize structures from this subset of data. In this report we will demonstrate the success of REDCRAFT in calculation of protein structures under these sparse data conditions.

REDCRAFT is also the only publicly available software package that is developed using a sound Object Oriented (OO) programming paradigm, and it therefore lends itself well to encapsulation of the physical and biophysical properties of proteins. For instance, the construction of a Polypeptide object from the more fundamental Atom and AminoAcid objects, directly reflects the natural process of polymerization and translates into better source code readability as well as faster development and program execution. In addition, OO design allows for easier extendibility of the system. For example, while the main data source of REDCRAFT is currently RDCs, one could easily extend the architecture to use NOEs or RCSAs. The only changes that the developer would need to make is the scoring mechanism of the elongation process and addition of any new atoms needed for the new data source. Existence of the AminoAcid class makes the addition of new atoms straightforward.

REDCRAFT’s approach to structure determination proceeds in two stages denoted as Stage-I and Stage-II. In Stage-I, a list of all possible torsion angles adjoining any two neighboring peptide planes is pruned and ranked based on structural fitness to the RDC data. Pruning of the local torsion angles can be based on scalar coupling data, maximum RDC fitness, dihedral constraints (such as Ramachandran or TALOS), or on evolutionary relationship to other proteins with an existing structure. Theoretically, the torsion angles adjoining any two peptide planes with the best fitness to the RDC data should constitute the correct geometry and therefore structure determination would be completed. Practically however, the globally optimal geometry will nearly always not be ranked as the first (due to experimental or structural noise); necessitating a more global search. Stage-II of REDCRAFT is designed to perform global optimization by elongating (similar to the elongation process during protein synthesis) a given fragment of size *N* peptide planes (initially a single dipeptide seed) by one peptide plane iteratively. Various conformers of the new extended fragment are systematically generated and ranked based on fitness to the RDC data (of the entire fragment). Typically, the top 2,000-10,000 structural candidates with the best fitness to the RDC data are propagated for extension in the next round of elongation. This process maintains a sufficiently diverse population of conformers to prevent entrapment in any one local minimum and is the primary reason for the success of REDCRAFT in characterization of protein structures with less data. The gradually increasing computational complexity of REDCRAFT is in stark contrast to the traditional methods of folding the entire protein at once. By starting the calculation for the entire protein, the algorithm must contend with the maximum level of complexity from the start, which transforms the problem into a global optimization. This, by in large, makes computation slower and limits the success due to inherent issues with high-dimensional global optimization problems such as entrapment in local minima. In contrast, REDCRAFT allows for structure calculation of a protein to proceed in an incrementally growing fragment that provides a optimization with a gradually increasing complexity. By utilizing this unique elongation approach, REDCRAFT is also capable of performing fragmented studies in which only certain sections of the protein can be characterized. Fragmented studies are useful in cases of missing or erroneous data. In the case of RDCs, which are very sensitive probes of internal dynamics, REDCRAFT’s fragmented study can be used to pinpoint the exact point of dynamics(34, 41, 70). It is also worth noting that even though the REDCRAFT engine only computes the backbone structure to reduce complexity, there are several methods available that can accurately calculate the sidechains of a protein based solely on the backbone(71).

REDCRAFT also provides several filtering and constraining tools that are uniquely useful for use with RDC data. For instance, Order Tensor Filter (OTF) allows selection of proteins based on prior knowledge of order tensors(72, 73). REDCRAFT also allows the user to define dihedral restraints. All restraints (including OTF and dihedral) can be flexibly turned on and off for select regions of a protein that may suffer from severe lack of experimental data. The most recent version of REDCRAFT (version 4.0) has also adopted NEF compliance in data import/export procedures, and has incorporated an advanced decimation process that has allowed for successful structure calculation of proteins with as much as ±4Hz of experimental noise(67, 69).

### Evaluation

Our evaluation of REDCRAFT was conducted in three phases with increasing level of difficulty in structure determination. In the first phase, REDCRAFT was tested using a set of proteins with existing experimental RDCs and X-ray or NMR structures. In the second phase, large proteins (larger than 500 residues) were chosen based on the availability of RDC data. Although a few large proteins have been subjected to RDC data acquisition, none contained enough RDC data to perform a meaningful structure calculation. In such instances, simulated RDCs were generated for a sparse set of interacting vectors. REDCRAFT was then used to calculate a RDC-based structure for each target protein to demonstrate the feasibility of RDC based structure calculation of larger proteins. The rationale for this phase is to illustrate the possibility of structure determination by RDCs when the collection of RDCs has been demonstrated in previous work. In the last phase of the study, a novel protein was targeted for a simultaneous study by RDC (using REDCRAFT) and NOE-based structure calculation. In each phase of the study structures calculated by REDCRAFT are compared to the existing NMR and X-ray structures (if applicable) of the respective proteins. The following sections provide more detailed information for each of the proteins as well as an overview of the REDCRAFT algorithm.

### Target Proteins

During the first phase of our experiment, we selected the target proteins (shown in Table 4) based on the availability of RDC data in BMRB or PDB, structural diversity, and existence of NMR or X-ray structure. RDC data for all the proteins except SR10 were obtained from the BMRB(30), while the RDC data for SR10 were obtained from the SPINE database(35). More detailed information regarding the exact RDCs can be found in the Table S1 of the Supplementary Material. Table 4 provides some self-explanatory information for each protein including the final column that highlights the average backbone similarity between the X-ray and NMR structures.

**Table 4.**
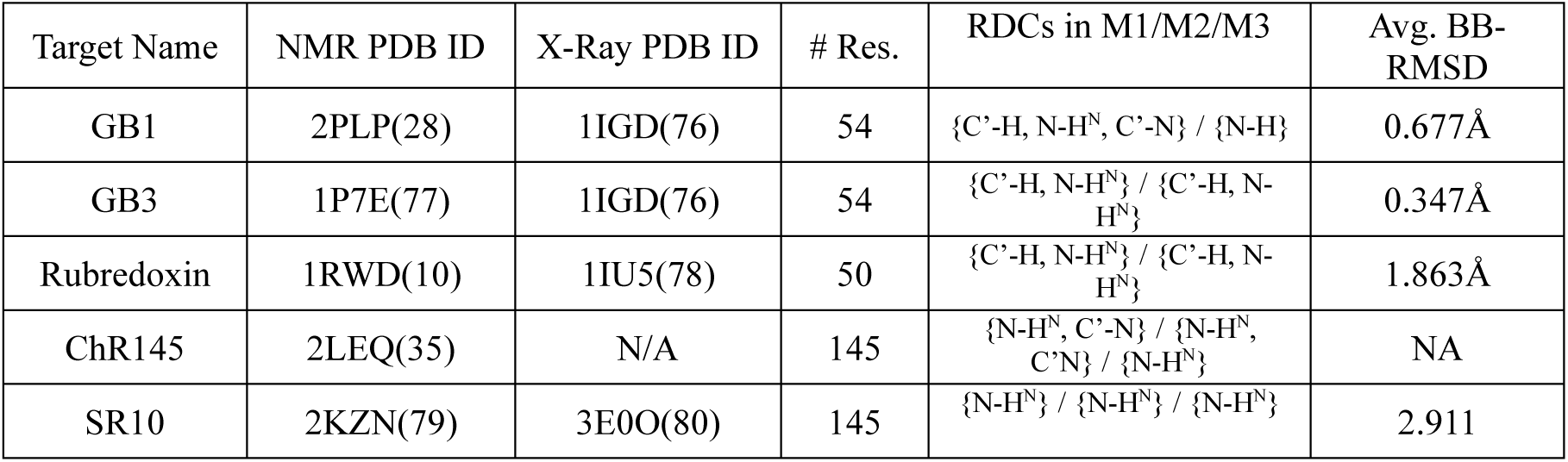
List of protein targets with their respective X-ray and NMR reference structures, RDCs used and the average BB-RMSD between the NMR and x-ray structures.

| Target Name | NMR PDB ID | X-Ray PDB ID | # Res. | RDCs in M1/M2/M3 | Avg. BB-RMSD |
| --- | --- | --- | --- | --- | --- |
| GB1 | 2PLP(28) | 1IGD(76) | 54 | {C'-H, N-H<sup>N</sup>, C'-N} / {N-H} | 0.677Å |
| GB3 | 1P7E(77) | 1IGD(76) | 54 | {C'-H, N-H<sup>N</sup>} / {C'-H, N-H<sup>N</sup>} | 0.347Å |
| Rubredoxin | 1RWD(10) | 1IU5(78) | 50 | {C'-H, N-H<sup>N</sup>} / {C'-H, N-H<sup>N</sup>} | 1.863Å |
| ChR145 | 2LEQ(35) | N/A | 145 | {N-H<sup>N</sup>, C'-N} / {N-H<sup>N</sup>, C'-N} / {N-H<sup>N</sup>} | NA |
| SR10 | 2KZN(79) | 3E0O(80) | 145 | {N-H<sup>N</sup>} / {N-H<sup>N</sup>} / {N-H<sup>N</sup>} | 2.911 |

The protein GB1 has been previously studied in depth(28, 74) and represents an ideal candidate to be used as a “proof of concept” case. GB3, an analog of GB1, was also investigated in this study using a different set of RDCs. The RDCs for the GB3 were previously collected(32, 33) for refinement of a solved crystal structure to obtain better fitness to experimental data (resulting in PDB ID 1P7E). Rubredoxin, represented another ideal target of study due to its mostly coil structure. Traditionally, structures that are heavily composed of helical regions prove difficult to solve for computational methods due to the near-parallel nature of their backbone N-H^N^ RDC vectors. ChR145, represents a larger, mixed beta-sheet and alpha helix protein. In a previous study, this protein was extracted from the Cytophaga hutchinsonii bacteria and characterized using traditional NMR restraints (primarily NOEs). Of interest, ChR145’s primary sequence is unique in the PDB. This fact alone makes its structural characterization difficult for any method that has a dependency on database lookups or homology modeling. SR10, a 145-residue protein, was characterized as part of the Protein Structure Initiative(75) and was included in this study to represent a challenging case because of the low RDC data density. The RDC data for this protein consisted of only {N-H^N^} vectors collected in three alignment media. Of additional interest, the RDCs were collected on a perdeuterated version of the SR10 protein.

Currently there are very few examples of large proteins in the BMRB database that include RDC data. If RDC data are available for large proteins, they are very scarce and from only one alignment medium. Meaningful structure determination of proteins from RDC data requires RDC data in two alignment media(81). Therefore, to investigate the feasibility of protein structure calculation of large proteins using only RDCs, synthetic sets of {C’-H, N-H^N^, C’-N}RDCs were generated in two alignment media using the software package REDCAT(61, 82) as described previously(83). A random error in the range of ±1 Hz was added to each vector to better simulate the experimental conditions. The proteins chosen for this controlled study are summarized in Table 5. Note that for ChR145 a synthetic study was also performed to demonstrate the unfragmented structure determination if additional RDC data had been acquired. It is noteworthy that Enzyme 1 from E. *coli* was chosen as an example of a large mixed α/β protein. The dataset used for solving the NMR structure of this protein included a very sparse set of {N-H^N^} RDCs that was not applicable in our studies, but demonstrates the possibility of RDC data collection in large proteins.

**Table 5.** List of protein targets used in the synthetic study of large proteins.

| Target Name | X-ray PDB ID | NMR PDB ID | # Res. | RDCs in M1/M2 |
| --- | --- | --- | --- | --- |
| ChR145 | 2LEQ(35) | N/A | 145 | {C'-H, N-H<sup>N</sup>, C'-N} / {C'-H, N-H<sup>N</sup>, C'-N} |
| Lpg1496 | 5T8C | N/A | 294 | {C'-H, N-H<sup>N</sup>, C'-N}/<br>{C'-H, N-H<sup>N</sup>, C'-N} |
| Enzyme 1 from *E. coli* | N/A | 2KX9(84) | 573 | {C'-H, N-H<sup>N</sup>, C'-N}/<br>{C'-H, N-H<sup>N</sup>, C'-N} |

In addition to the previously characterized proteins, RDC data were acquired for a novel, 71-residue protein (designated PF2048.1). PF2048.1 has been selected as a target of our studies due to its novelty in comparison to the existing archive of structurally characterized proteins. PF2048.1, an all-helical 9.16 kDa protein, exhibited less than 12% sequence identity to any structurally characterized protein in PDB (as of January 2019). The previously reported computational models of this structure(72) concluded the helical nature of this protein and resulted in an ensemble of structures with as much as 10Å of backbone diversity(68, 72, 73).

RDC data were acquired by NMR spectroscopy for this protein in Phage and stretched Poly Acrylamide Gel (PAG) alignment media. The resulting two sets of RDCs consisted of {N-C’, N-H^N^, C’-H} from the Phage and {N-H^N^} from the PAG media. The process of NMR data collection is described at length in Section 2.3. Collectively, the two data sets were missing ∼17% of data points (48/276) leaving 228 total RDC data points (an average of 1.6 RDCs per residue, per alignment medium).

### PF2048.1

#### Expression and Purification

Expression and purification were performed by Nexomics Inc. Prior to gene synthesis, the sequence was optimized by codon optimization software. The designed gene was synthesized by Synbio-Tech (www.Synbio-tech.com) and subcloned into pET21-NESG vector.

Protein expression was performed as previously reported(85). Briefly, the recombinant pET21-NxSC1 plasmid was transformed into *E. coli* BL21 (DE3) cells and the cells were cultured in ^13^C^15^N-MJ9 medium containing 100 μg/mL of ampicillin. The culture was further incubated at 37°C and protein expression was induced by addition of isopropyl β-D-1-thiogalactopyranoside (IPTG) to the final concentration of 1 mM at logarithmic phase. Cells were harvested after overnight culture at 18°C and protein expression was evaluated by SDS-PAGE.

The protein was purified using a standard Ni affinity followed by size exclusion two-step chromatography method first as previously reported(85). Since the purified NxSC1 sample presented as 2 bands, an additional ion exchange chromatography was performed. The NxSC1 sample from the two-step purification was pooled and dialyzed against buffer A (Buffer A: 20 mM Tris-HCl, pH 7.5), and loaded onto a HiTrap Q HP 5 ml column. A gradient of NaCl from 0 to 1 M was applied (Buffer B: 20 mM Tris-HCl, pH 7.5, 1 M NaCl). The NxSC1 was pooled and concentrated to 1 mM using Amico Ultra-4 (Millipore).

### NMR Sample Preparation and Data Acquisition of PF2048.1

For measurements under isotropic conditions a sample of PF2048.1 was prepared at a concentration of 0.8 mM in 20 mM MES, 100 mM NaCl, and 5 mM CaCl_2_ at pH 6.5. All samples also contained 10 mM DTT, 0.02% NaN_3_, 1 mM DSS, and 10% D2O. An anisotropic sample is required for the measurement of RDCs. After isotropic data collection, the PF2048.1 sample was used to prepare two partially aligned samples. A sample with pf1 phage as the alignment medium (designated alignment medium M1) was prepared which contained 0.88 mM PF2048.1 and 48 mg/mL phage in Tris buffer. After equilibration at room temperature for 10 min at 25 °C the sample showed a deuterium splitting of 8.8 Hz when placed in the magnet. A second aligned sample was prepared in a 5 mm Shigemi tube using positively charged poly-acrylamide compressed gels (designated alignment medium M3). This sample contained approximately 0.77 mM PF2048.1. After equilibration at 4 °C for 7–8 h the sample showed uniform swelling of the gel which is compressed vertically.

NMR data were collected at 25°C using Bruker Avance II 600 and 800 MHz spectrometers equipped with 5-mm cryoprobes. Sequence specific backbone and side-chain NMR resonance assignments were determined using standard triple-resonance NMR experiments (Table S2). Processing of NMR spectra was done using TopSpin and NMRPipe whereas visualization was done using NMRDraw and Sparky. NMR spectra were analyzed by consensus automated backbone assignment analysis using PINE(86) and AutoAssign(87) software, and then extended by manual analysis to determine resonance assignments. The solution NMR stricture was calculated using CYANA using backbone and side-chain chemical shifts, ^15^N and^13^C-edited NOESY, and phi (*φ) and psi* (*Ψ)* dihedral angle constraints derived from TALOS. Calculations using CYANA were done iteratively to refine NOESY peak lists, verify and complete resonance assignments using interactive spectral analysis with RPF(88) and Sparky(86) software. The 20 lowest energy conformers of 100 calculated structure, based on target function scoe, were further refined by simulated annealing and molecular dynamics in explicit water using CNS. Structure quality scores were calculated using Protein Structure Validation Suite (PSVS)(89) and the goodness-of-fit between the NOESY peak lists and the final ensemble of conformers were derived from RFP software.

### Structure Calculation with NOEs

Uniformly ^13^C,^15^N-enriched PF2048.1, a 72-residue protein, was used for NMR structure determination, using standard triple-resonance NMR methods outlined in the Methods and Materials. The structure was determined from ^15^N-and ^13^C-resolved 3D-NOESY data, both with and without RDC restraints. In total 217 RDC measurements were used: 54 HC’, 54 NC’, 57 HN RDC from medium 1 (M1 – phage), and 52 HN RDC from medium 2 (M3 –stretched polyacrylamide gel). Both ASDP(90) and CYANA3.97(17) were used to automatically assign long-range NOEs and to calculate these structures. ASDP(90) was also used to guide the iterative cycles of noise/artifact NOESY peak removal, peak picking and NOE assignments, as described elsewhere(91). NOE matching tolerances of 0.030, 0.03 and 0.40 ppm were used for indirect ^1^H, direct ^1^H, and heavy atom ^13^C/^15^N dimensions, respectively, throughout the CYANA and ASDP calculations. This analysis provided > 2,300 NOE-derived conformationally-restraining distance restraints (Supplementary Tables S1 and S2). In addition, 132 backbone dihedral angle restraints were derived from chemical shifts, using the program TALOS_N(92), together with 70-74 hydrogen-bond restraints. Structure calculations were then carried out using ∼ 35 conformational restraints per residue. One hundred random structures were generated and annealed using 10,000 steps. Similar results were obtained using both Cyana and ASDP automated analysis software programs. The 20 conformers with the lowest target function value from the CYANA calculations were then refined in an ‘explicit water bath’ using the program CNS and the PARAM19 force field(93), using the final NOE derived distance restraints, TALOS_N dihedral angle restraints, and hydrogen bond restraints derived from CYANA. Structure quality factors were assessed using the PDBStat(88) and PSVS 1.5(89) software packages. The global goodness-of-fit of the final structure ensemble with the NOESY peak list data were determined using the RPF analysis program(88).

### Structure Calculation with REDCRAFT

REDCRAFT(13, 34, 40, 41, 67, 68, 70, 94) was used to calculate the structure of proteins from RDC data with a standard depth search of 1000. Additional features such as decimation(41), minimization(34), and 4-bond LJ(34) terms were included in all calculations. Other features such as Order Tensor Filter(34), and Dynamically Adaptive Decimation(69) were not used in this exercise. For evaluation purposes, the RDC-RMSD reported by REDCRAFT was converted to Q-factor to assess the final models’ fitness to RDC data. The backbone-RMSD (BB-RMSD) of REDCRAFT structures to existing structures were calculated using the *align* function of PyMOL(95) without the exclusion of any atoms.

Under certain circumstances, structure determination by REDCRAFT is recommended to be conducted in discrete fragments. One such instance is based on gap in the experimental data. In comparison to the NOE-based structure determination, this can be a very powerful feature. Fragmented study of a protein allows direct study of a certain region of interest in a protein and therefore reduce the overall cost of data acquisition. A second instance relates to structure determination of large proteins, during which accumulation of structural noise may influence the course of structure determination. Departure from ideal peptide geometries (e.g. planarity of the peptide plane), variation in bond distances, and bond angles can be cited as examples of structural noise. The effect of structural noise, sometimes, accrues to become noticeable for fragments larger than one hundred amino acids. Any existing gaps of less than 6 amino acids can easily be filled during the process of structural refinement. In this study we have used XPLOR-NIH(15) to address variation from ideal peptide geometries and complete the missing gaps. More specifically, during the final refinement process, each structure was subjected to 30,000 steps of Powell minimization that included the same set of RDCs used during the structure calculation with REDCRAFT. Aside from completion of the missing residues, these minimizations normally resulted in structural variation of less than 0.5A.

## Acknowledgements

Funding for this project was provided by NIH grants P20 RR-016461 (to HV) and 1R01GM120574 (to GTM).

## Supplementary Information

S1 Table. Structure Calculation Input Files

S2 Table. Structure Quality Statistics

S1 Figure. Solution NMR structure of PF2048.1 refined with RDC (green) and without RDC (cyan). The overall RMSD is within 0.6 Å.

